# Phosphatase-independent suppression of mucosal inflammation and disease progression by *Salmonella* SopB

**DOI:** 10.1101/2024.09.02.610793

**Authors:** Nour Diab, Eva-Lena Stange, Chiun Huei Yong, Jörg Deiwick, Mihael Vucur, Tom Lüdde, Michael Hensel, Natalia Torow, Kaiyi Zhang, Mathias W. Hornef

## Abstract

*Salmonella enterica* subsp. *enterica* serovar Typhimurium (*S.* Typhimurium) translocates effector molecules via its *Salmonella* pathogenicity island (SPI)1 encoded type 3 secretion system (T3SS) to induce internalization by intestinal epithelial cells and manipulate cellular responses. Among these effector molecules, the *Salmonella* outer protein B (SopB) was shown to possess phosphatidyl-inositol phosphatase activity and induce bacterial internalisation, promote cell survival, influence endosomal trafficking and alter host cell signalling. Using a neonatal *S.* Typhimurium infection model, we here show that SopB *in vivo* suppresses early epithelial chemokine expression, delays mucosal immune cell recruitment, reduces barrier impairment by enterocyte necroptosis, and prevents disease progression and premature death. Unexpectedly, this immunosuppressive effect was independent of the phosphatidyl-inositol phosphatase and phosphotransferase activity of SopB but required an intact N-terminal domain. Thus, SopB exerts a potent phosphatase-independent immunosuppressive effect to delay local tissue inflammation and disease progression likely to promote host transmission.

## Introduction

The human enteropathogen *Salmonella enterica* subsp. *enterica* sv. Typhimurium (*S.* Typhimurium) uses the *Salmonella* pathogenicity island (SPI)1-encoded type three secretion system (T3SS) to translocate effector molecules into intestinal epithelial cells. These effector molecules induce internalisation, shape formation of the *Salmonella* containing vacuole (SCV) and manipulate host cell processes to promote intracellular survival and proliferation ^1–3^. Among the effector molecules translocated by the SPI-1 T3SS is SopB, a phosphatidyl-inositol phosphate 4 and 5 (PtdIns(4,5)P_2_) phosphatase and phospho-transferase/isomerase. SopB generates PtdIns(3)P_1_, PtdIns(3,4,5)P_3_ and PtdIns(3,4)P_2_ in a non-canonical phosphoinositide 3-kinase (PI3K)-independent manner ^4–7^. SopB is ubiquitously present in human and animal *Salmonella* isolates ^8^.

Through its C-terminally encoded phosphatase domain (aa 357-561), SopB together with SopE/SopE_2_ activates the Rho GTPases Cdc42 ^9^ and enriches RhoJ, RhoB, RhoH and R-Ras1 ^10^, PtdIns(3,4)P_2_ and PtdIns(3,4,5)P_3_ ^11^, Arf GEF ARNO ^12^, Annexin A2 ^13,14^, Myo6 ^15^, as well as SNX9 ^7^ and SNX18 ^16^ at the cell entry site, leading to actin remodelling, ruffling, membrane fission and uptake by intestinal epithelial cells, the first step in mucosal translocation and invasive infection ^5,17^. SopB is then multi-mono-ubiqutinated, recruited to the endosomal membrane and reduces the negative surface charge of the SCV by removing PtdlIns(4,5)P_2_ and phophatidyl-serine, thereby inhibiting Rab recruitment and lysosomal fusion ^18–21^. Additionally, SopB induces a sustained phosphatase-dependent Akt phosphorylation promoting cell survival ^22–28^. SopB expression persists for many hours *in vivo* ^25,29^.

More recently, the N-terminal GTPase binding domain (residues 117-168) of SopB has been shown to facilitate phosphatase-independent interaction with Cdc42 leading to actin fibre disruption, cell cycle arrest and MAP kinase signalling in yeast cells ^30–34^. The functional role of this N-terminal domain in eukaryotic cells and during *in vivo* infection has not systematically been analyzed.

Using our oral neonatal mouse infection model we have previously shown, that SopB *in vivo* is not required for enterocyte invasion or the intraepithelial formation of SCVs ^3^. Here, we analysed the role of SopB during the early course of the infection using isogenic bacterial mutants and mice deficient in innate immune signalling and cell death pathways. Unexpectedly, we observed accelerated disease progression and early mortality of neonatal mice after infection with SopB mutant *S.* Typhimurium. This phenotype was independent on the phosphatidyl-inositol phosphatase activity of SopB but associated with an early increase in epithelial chemokine expression, mucosal immune cell recruitment, and enterocyte necroptosis.

## Material and Methods

### Bacterial strains

In this study, wildtype (wt) *Salmonella enterica* subsp. *enterica* serovar Typhimurium (ATCC14028, *S*. Typhimurium), an isogenic Δ*sopB* mutant strain, Δ*sopB* mutant strains complemented with *sopB* (Δ*sopB,* p*sopB*) or *sopB* and its chaperon *sigE* (Δ*sopB,* p*sopBsigE*), a triple *sopB*, *sopE_2_*, *sipA* (Δ*sopBE_2_sipA*) and sopA, sopE2, sipA (Δ*sopAE_2_sipA*) strain, as well as strains carrying a chromosomal point mutation in SopB in the C-terminal domain at position 460 (C460S, SopB*^C^*^460^*^S^*) and 528 (K528A, SopB*^K^*^528^*^A^*) or in the N-terminal domain in position 76 (L76P, SopB*^L76P^*) were used. For *in vitro* experiments, wt and Δ*invC S*. Typhimurium carrying p4042 encoding *sopB*::HA, the triple mutant Δ*sopAE_2_sipA S*. Typhimurium encoding a chromosomal *sopB*::HA fusion protein and the quadruple mutant Δ*sopABE_2_sipA* carrying p4042 encoding *sopB*::HA were used.

### Ethics statement

All animal experiments were performed in compliance with the German animal protection law (TierSchG) and approved by the local animal welfare committee (Niedersachsische Landesamt für Verbraucherschutz und Lebensmittelsicherheit Oldenburg, Germany; Landesamt für Natur, Umwelt und Verbraucherschutz, North Rhine Westfalia) under the code 8–02.04.2021.A043 and 40152A4 including all approved changes.

### *In vivo* infection experiments

Adult C57BL/6J wild type mice, *MyD88^flox/flox^* mice (B6.129P2(SJL)-Myd88tm1Defr/J; stock 008888) crossed with vav*-Cre^+^*(B6.Cg-Commd10Tg(Vav1-icre)A2Kio/J; stock 008610) animals, *Casp1*^-/-^ (B6. 129S2-Casp1^tm1Flv^/J, stock no.016621) and *Tnfr1*^-/-^ (Tnfrsf1a^tm1MAK^; stock 002818) were obtained from Jackson Laboratory (Bar Harbour, USA) and bred locally at University Hospital RWTH Aachen under SPF conditions. *Asc*^-/-^ mice (B6. 129-Pycard^tm1Vmd^), *Mlkl*^-/-^ (BV6. 129-Mlkl^tm1/J^), and *Casp8^ΔIEC^* (B6. 129-Casp8^tm1Hed^/J; stock 027002) mice were provided by James Murphy (Walter and Eliza Hall Institute of Medical Research, Australia) and Claudia Günther (University Hospital Erlangen, Germany) and bred locally at University Hospital RWTH Aachen under SPF conditions. Overnight bacterial cultures were diluted 1:10 and incubated at 37°C until reaching the logarithmic phase (OD_600_: 0.5). Bacteria were diluted to obtain the appropriate inoculum in PBS. One-day-old animals were orally infected with 100 CFU *S.* Typhimurium. At the indicated time point postinfection (p.i.), liver, lung, spleen and mesenteric lymph nodes (MLN), small intestine as well as blood samples were collected. Viable counts were obtained by serial dilution and plating of homogenized tissue on LB agar plates supplemented with the appropriate antibiotic(s). Small intestinal tissues were collected, fixed in 4% PFA or snap frozen in liquid nitrogen for histological analysis and total tissue expression.

### Gene expression analysis

RNA was isolated from the epithelial cell pellet or homogenized intestinal tissue using TRIzol® according to manufacturer’s recommendations. The RNA concentration was determined using a NanoDrop 1000 spectrophotometer (Thermo Scientific). First-strand complementary DNA (cDNA) was synthesized from 5 µg total RNA using Oligo-dT primers, 5X PCR buffer, dNTP, RevertAid reverse transcriptase and RiboLock RNAse inhibitor (ThermoFisher Scientific). RT-PCR was Taqman technology with an absolute QPCR ROX mix (Thermo Scientific). Taqman probes *Hprt* (housekeeping gene, Mm00446968-m1), *Cxcl1* (Mm04207460_m1), *Cxcl2* (Mm00436450_m1), *Cxcl5* (Mm00436451_g1), *Mcp1* (Mm00441242_m1), *Tnf-α* (Mm00443258_m1), *Reg3γ* (Mm00441128_g1), *Bcl2* (Mm01302952_m1) or *iNos* (Mm00440502_m1) from ThermoFisher Scientific were used. Results were calculated by the 2^−ΔΔCt^ method. Values were normalized to the *Hprt* housekeeping gene and are presented as fold induction over age-matched healthy controls.

### Cytokine and chemokine quantification

Serum cytokine and chemokine levels were measured using the Cytometric Bead Array Kit (BioLegend) according to the manufacturer’s protocol. Samples were analyzed with a BD FACS Canto II and analyzed with FlowJo X. Results are expressed as picograms per milliliter. The heat map was generated using Heatmapper.

### Intestinal epithelial cell and immune cells isolation and analysis

Primary small intestinal epithelial cells were isolated as previously described ^2^. Briefly, epithelial cells were detached from the underlying tissue in 30 mM EDTA/PBS with strong shaking. To analyze the number of intraepithelial *S.* Typhimurium, one fraction of epithelial cells was treated with 100 μg/mL gentamicin for 1 h at room temperature, washed in PBS and plated in serial dilutions on selective LB agar plates. The other fraction of epithelial cells was stored at -80°C for subsequent gene expression analysis. For the isolation of immune cells, the *lamina propria* was enzymatically digested in RPMI (Gibco) containing 30 µg/ml liberase^TM^ (Roche) and 50 µg/ml DNAse (Roche) for 45min shaking at 37°C. Tissue pieces were filtered through a 100 µm nylon cell strainer (BD) to obtain a single cell suspension. Cells were stained using the following antibodies CD45-BV510, CD45-PE Cy7, Ly6C-PerCPCy5.5, Ly6G-PE, Ly6G-Spark NIR 685, Ly6C-BV711, CD11b-APC Cy7, CD11b-BUV 395, CD64-APC, CD64-PE Dazzle, MHCII-AF488, MHCII-BV510, PDL1-PE, SiglecF-APC R700, Epcam-BV421, CD3-FITC, CD19-FITC, and DAPI (Roth) for subsequent analytical flow cytometry. Data were acquired with a BD FACS Canto II and analyzed with FlowJo X. For FACS sorting, approximately 3-6 million monocytes, macrophages, neutrophils and eosinophils were sorted by BD Biosciences FACS Aria Fusion Sorter and directly collected into RNA lysis Buffer (QIAGEN RNeasy Micro Kit QIAGEN). Total RNA of each immune cell population was extracted using QIAGEN RNeasy Micro Kit following manufacturer’s instructions.

### *Ex vivo* stimulation of isolated immune cells

Immune cells were isolated and separated using a Percoll gradient by centrifugation at 700 x g for 20 min. at room temperature. Cells were collected, washed in 3% FCS/PBS and counted. 10^6^ cells were cultured in 500 µL Iscove’s modified Dulbecco’s medium (Thermo Fisher Scientific) supplemented with 10% FBS, with/without 1 µL of the cell activation cocktail (phorbol myristate acetate (PMA) and Ionomycin (I), Biolegend, Cat. No.: 423302). After incubating the cells at 37°C in a 5% CO_2_ humidified atmosphere for 1 h, 5 µg/ml brefeldin A (BFA, Biolegend, Cat. No.: 420601) was added. After 3 h of restimulation, cells were collected, washed once with FACS buffer, and resuspended in 3% FCS/PBS. Cells were then harvested and stained with the following antibodies: CD45-APC R700, CD3-APC Fire750, PDL1-APC, SiglecF-BB 515, CD11c-BUV 737, CD64-PE Dazzle, CD11b-BV 786, F480-PE Cy5, Epcam-BV421, MHCII-BV510, Ly6C-PerCP Cy5.5, Ly6G-BV711, CD80-BUV 805, CD19-PE Cy7 and Zombie UV (Cat. No.: 423107, BioLegend). Stained cells were fixed and permeabilized prior to intracellular cytokine staining with a TNFα-PE mouse antibody (Biolegend). Data were acquired on a Cytek Aurora flow cytometer and analyzed with FlowJo X.

### Immunostaining

Small intestine tissues were fixed in 4% paraformaldehyde (PFA) and embedded in paraffin blocks. 4 μm thick sections were deparaffinized and rehydrated. Slides were stained with hematoxylin and eosin Y solution for H & E staining and observed under a Zeiss Axoi Imager M2 light microscope. The thickness of the *lamina propria* as a measure of the tissue edema was measured using the ZEN 3.4 imaging software. For immunofluorescence staining, tissue sections were blocked with 10% normal donkey serum/5% BSA/PBS. Rabbit anti-Ki67 (Ab15580, Abcam), mouse anti-E-cadherin (610182, BD Transduction Laboratories), rat anti-PMN (Ly6-6B2, SeroTec), rabbit anti-Muc2 (GTX100664, BIOZOL) were used at the appropriate dilution followed by the fluorophore-conjugated donkey secondary antibody (Jackson ImmunoResearch). AF647-conjugated wheat germ agglutinin (WGA, Vector Laboratories, FL1021) was used to visualize the mucus layer. The *in situ* cell death detection Kit (Roche) was used following manufacturer’s instructions to detect TUNEL positive cells. Slides were counterstained with DAPI (Vector Laboratories) and images were taken using a Zeiss ApoTome.2 system microscope connected to an Axiocam 506 digital camera (Zeiss).

### Neonatal intestinal epithelial stem cell organoid culture

Neonatal intestinal epithelial stem cell organoids (spheroids) were prepared according to established protocols and grown as cell monolayers ^35,36^. R-spondin producing HEK293T cells and Wnt3a producing HEK23T cells were kindly provided by Calvin Kuo (Stanford University, Stanford, CA, USA) and Sina Bartfeld (Berlin Technical University, Berlin, Germany), respectively. Confluent cell monolayers were infected with *S*. Typhimurium at a multiplicity of infection (MOI) of 10:1 for 1 h. Monolayers were washed three times with warm PBS and supplemented with pre-warmed ENRWY media containing 100 μg/mL gentamicin (Sigma) for 1h at 37°C to remove extracellular bacteria. Infected monolayers were washed again three times in warm PBS and lysed. The number of intracellular bacteria was determined by serial dilution and plating on selective LB agar plates. The invasion rate was calculated as (number of intracellular bacteria/number of administered bacteria) × 100[%].

### Programmed cell death assay using L929 cells

L929 cells were cultured in Dulbecco’s modified Eagle’s medium (PAN BIOTECH) supplemented with 10% heat-inactivated FCS until confluency. Confluent cells were pretreated for 1 h with DMSO only, Z-vad (5 µM, Pan-caspase inhibitor, R&D systems), Z-vad plus SopB (3 µg/mL, provided by Jörg Deiwick and Michael Hensel, Osnabrück, Germany) or Z-vad plus necrostatin-1s (10 µM, Nec-1s, Selleckchem, Cat. No.: S8641). After 1 h, the supernatant was discarded the cells were incubated for 20 h in medium supplemented with DMSO, Z-vad, Z-vad with TNF-α (10 ng/mL, GenScript), Z-vad plus SopB and TNF-α, Z-vad plus Nec-1s, or Z-vad plus SopB and Nec-1s. To induce extrinsic apoptosis, cells were treated with DMSO only or preincubated for 1 h with CHX (10 µg/ml, Sigma-Aldrich, Cat. No.: C6255) plus Nec-1s (10 µM). Then, the supernatant was removed, and the cells were treated with TNF-α (10 ng/ml), CHX and Nec-1s for 15 h to induce apoptosis. To block apoptosis, cells were preincubated with SopB (0.1 µg/ml), CHX and Nec-1s or Z-vad, CHX and Nec-1s for 1 h and then 10 ng/ml TNF-α was added to the cells. 15 and 20 h after stimulation, respectively, apoptotic and necroptotic cells were quantified under bright field of ZEISS Apotome.2 system microscope for morphological changes. Images were analyzed using the ZEN 3.4 imaging software. For flow cytometry analysis, the L929 cells were harvested at 300 x g for 4 min at 4°C, washed twice with cold FACS buffer (3%FCS/PBS) and analyzed for apoptosis or necroptosis using the APC Annexin V Apoptosis Detection Kit with propidium iodide (PI) (Biolegend) following the manufacturer’s instructions. Data were acquired with a BD FACS Canto II and analyzed with FlowJo X.

### Western Blot

Total protein was precipitated from the cell-free supernatant of a mid-log *S. Typhimurium* culture using trichloroacetic acid (TCA). *S. Typhimurium* cells were collected by centrifugation and lysed (bacterial pellet). Following electrophoresis in a 10% acrylamide gel, immunoblotting was performed and HA-tagged SopB was detected using a HA-specific antibody (diluted 1:1000, Roche, Cat. No.: 1867423).

### Statistics

Mortality was analyzed by log-rank (Mantel-Cox) test. The non-parametric Mann Whitney test was used for the comparative analysis of two groups; the One-way ANOVA with Kruskal-Wallis, Bonferroni or Tukey’s multiple comparison test was employed for the statistical analysis of more than two groups. Two-way ANOVA with Sidak or Tukey’s multiple comparison test was employed for the statistical analysis of two groups that have been split on two independent variables. Graphpad Prism Software 9 was used for statistical evaluation. Differences were considered significant at p <0.05, *; p <0.01, **; p <0.001, ***; and p <0.0001 (****).

## Results

### Accelerated disease progression in the absence of SopB

Infection of newborn mice with a *Salmonella enterica* subsp. *enterica* sv. Typhimurium (*S.* Typhimurium) strain lacking the *Salmonella* pathogenicity island (SPI)1-associated phosphatidyl-inositol phosphatase SopB (Δ*sopB*) resulted in significantly earlier mortality as compared to wildtype (wt) *S.* Typhimurium infection (Fig. 1A). Complementation (compl.) with an expression plasmid carrying *sopB* together with its chaperone *sigE* reversed this phenotype. SopB has previously been reported to contribute to intestinal epithelial cell invasion *in vitro.* Consistently, Δ*sopB S.* Typhimurium showed moderately reduced cell invasion in an intestinal epithelial stem cell organoid monolayer co-culture model *in vitro* (Suppl. Fig. 1A and B) ^7,9,12,21^. Following *in vivo* infection, however, no difference in the number of intraepithelial bacteria (Fig. 1B) or bacterial counts in total mesenteric lymph node, spleen, and liver tissue between Δ*sopB* and wt *S.* Typhimurium was observed at day 1 and 2 p.i. (Fig. 1C and D). These results suggested that the SPI1 effector SopB *in vivo* controls disease progression and reduces early infection-induced mortality. This *in vivo* phenotype most likely does not result from altered enterocyte invasion, intracellular survival and proliferation, or tissue spread.

**Figure 1.**
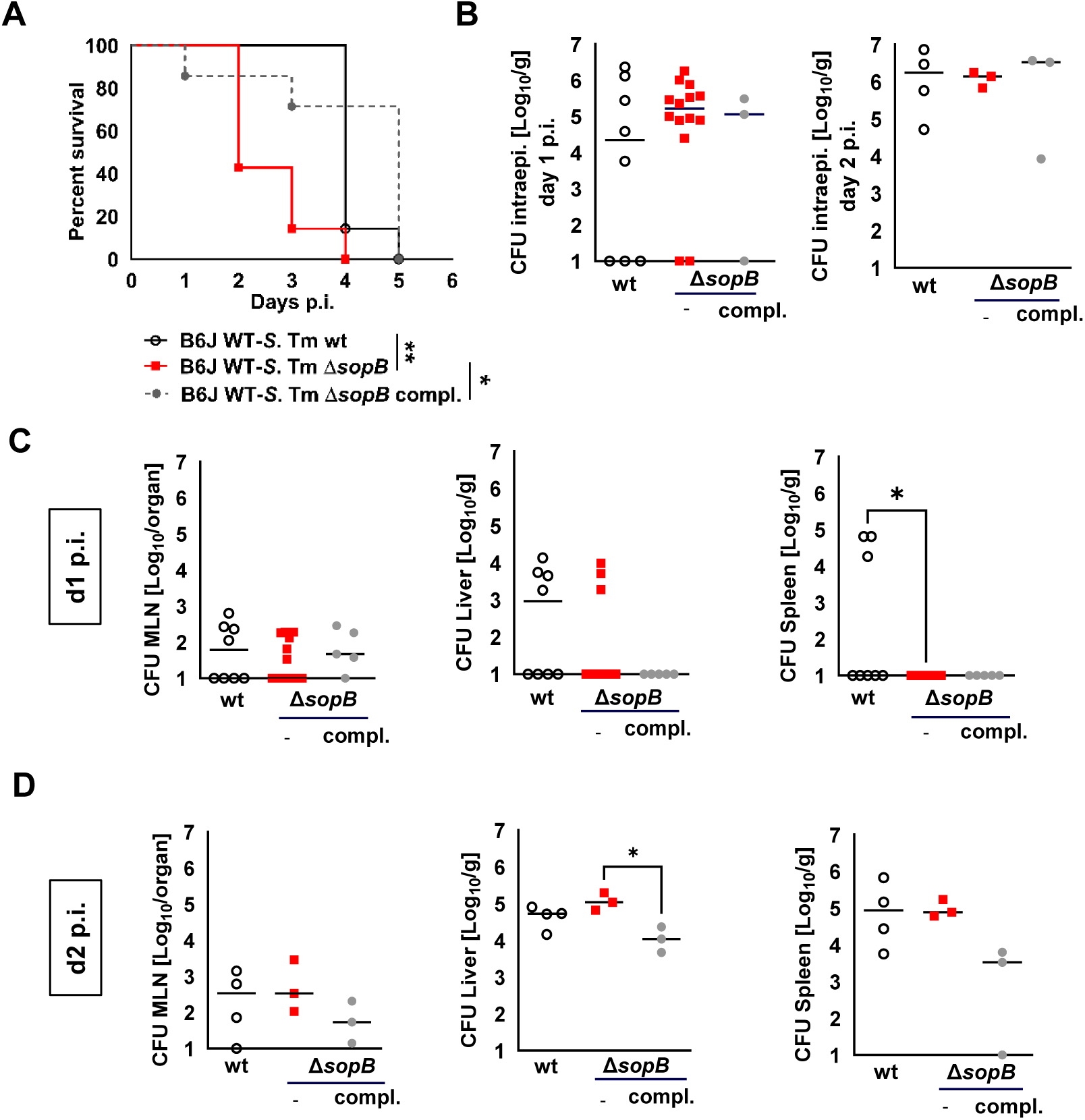
Survival and bacterial organ load. **(A)** Kaplan-Meier curve of 1-day-old C57BL/6 (B6JWT) neonates orally infected with 100 CFU wt (n=7), *sopB* deficient (Δ*sopB*, n=7), or *sopB* deficient *S*. Typhimurium complemented in trans with *sopB* and its chaperon sigE (Δ*sopB* p*sopBsigE*, compl., n=7). log-rank (Mantel-Cox) test. **(B-D)** Number of intracellular *S.* Typhimurium in isolated intestinal epithelial cells at day 1 and 2 p.i. **(B)** and total bacterial count in mesenteric lymph node (MLN), spleen and liver tissue at day 1 **(B)** and 2 **(C)** post infection (p.i.). One data point represents one animal, mean, n=3-14 animals per group from at least two independent experiments. One way ANOVA. *, p<0.05; **, p<0.01.

### SopB dampens the early epithelial host response and mucosal tissue damage

Intestinal epithelial expression of the chemokines *Cxcl2*, *Cxcl1* and *Mcp1* was markedly increased in neonatal animals infected with Δ*sopB* as compared to wt *S.* Typhimurium at day 1 and 2 p.i (Fig. 2A and B). In fact, Δ*sopB* infected neonates exhibited a strongly enhanced epithelial chemokine expression already at day 1 p.i., whereas this increase was completely blunted after wt infection. Also, epithelial expression of the antimicrobial protein *Reg3γ* was increased at day 1 and 2 and the chemokine *Cxcl5* was increased at day 2 p.i. with Δ*sopB* as compared to wt *S.* Typhimurium (Suppl. Fig. 2A and B). Serum levels of the cytokines IL-1β, IFN-γ and IP10 and the chemokine Ccl5 were significantly higher in Δ*sopB* as compared to wt *S.* Typhimurium-infected mice at day 2 p.i. (Fig. 2C and Suppl. Fig. 2C-O). Consistent with increased early chemokine expression, the number of monocytes and neutrophilic granulocytes but not macrophages in the mucosal small intestinal tissue was strongly enhanced in Δ*sopB S.* Typhimurium-infected mice at day 2 p.i. (Fig. 3A-E and Suppl. Fig. 3). Histological analysis further revealed significantly enhanced thickening of the submucosal tissue indicating edema formation (Fig. 4A and B), goblet cell depletion, reduced goblet cell size suggesting enhanced mucus secretion (Fig. 4C-E), and elevated numbers of TUNEL positive enterocytes indicating reduced epithelial integrity (Fig. 4F and G) after Δ*sopB* as compared to wt *S.* Typhimurium infection. In addition, Δ*sopB S.* Typhimurium-infected mice exhibited increased intestinal epithelial cell proliferation illustrated by elevated numbers of Ki67-positive cells suggesting epithelial repair (Fig. 4H and I). Thus, SopB controls the early mucosal host response, immune cell infiltration and tissue damage upon *S.* Typhimurium infection

**Figure 2.**
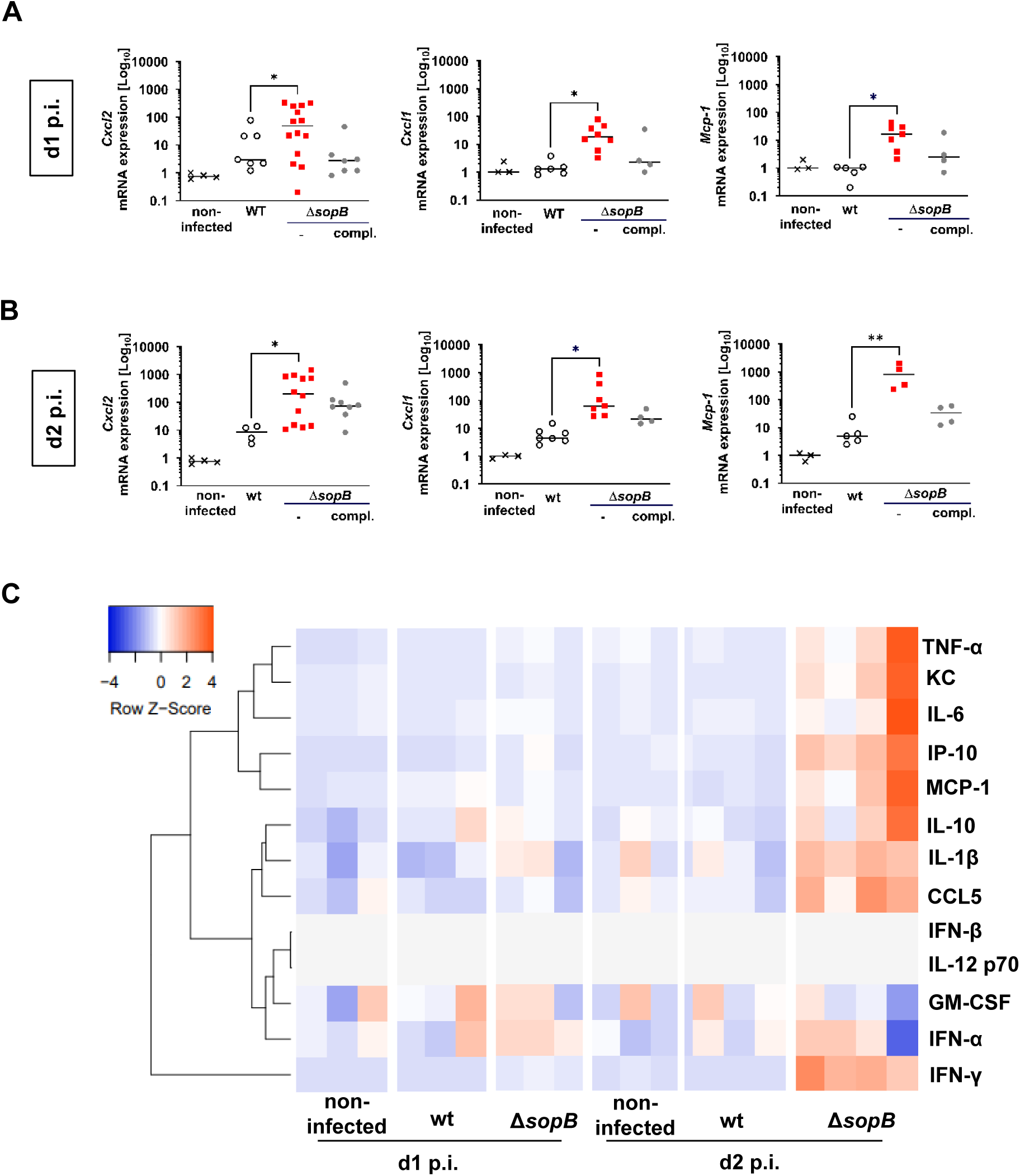
Chemokine and cytokine expression. **(A and B)** 1-day-old C57BL/6 neonates were left untreated (n=3-4) or orally infected with 100 CFU wt (n=4-7), *sopB* deficient (Δ*sopB*, n=4-14), or *sopB* deficient *S*. Typhimurium complemented in trans with *sopB* and its chaperon *sigE* (Δ*sopB* p*sopBsigE*, compl., n=4-8). *Cxcl1*, *Cxcl2* and *Mcp1* expression in total isolated intestinal epithelial cells at day 1 **(A)** and day 2 **(B)** p.i.. Values were normalized to the house keeping gene *Hprt* and are showed as fold expression over uninfected age-matched control animals. One data point represents one animal from at least two independent experiments, median. One-way ANOVA with Kruskal-Wallis comparison test. **(C)** Color-scaled heat map (z-score) showing the concentration of the indicated cytokines and chemokines in the serum of uninfected age-matched animals (n=3), or mice infected at day 1 after birth with 100 CFU wt (n=3) or *sopB* deficient *S*. Typhimurium (Δ*sopB*, n=4) at day 1 and 2 p.i. Each column represents one animal. Mean ± SD, two-way ANOVA with Tukey’s multiple comparison test. One data point represents one animal. *, p<0.05; **, p<0.01.

**Figure 3.**
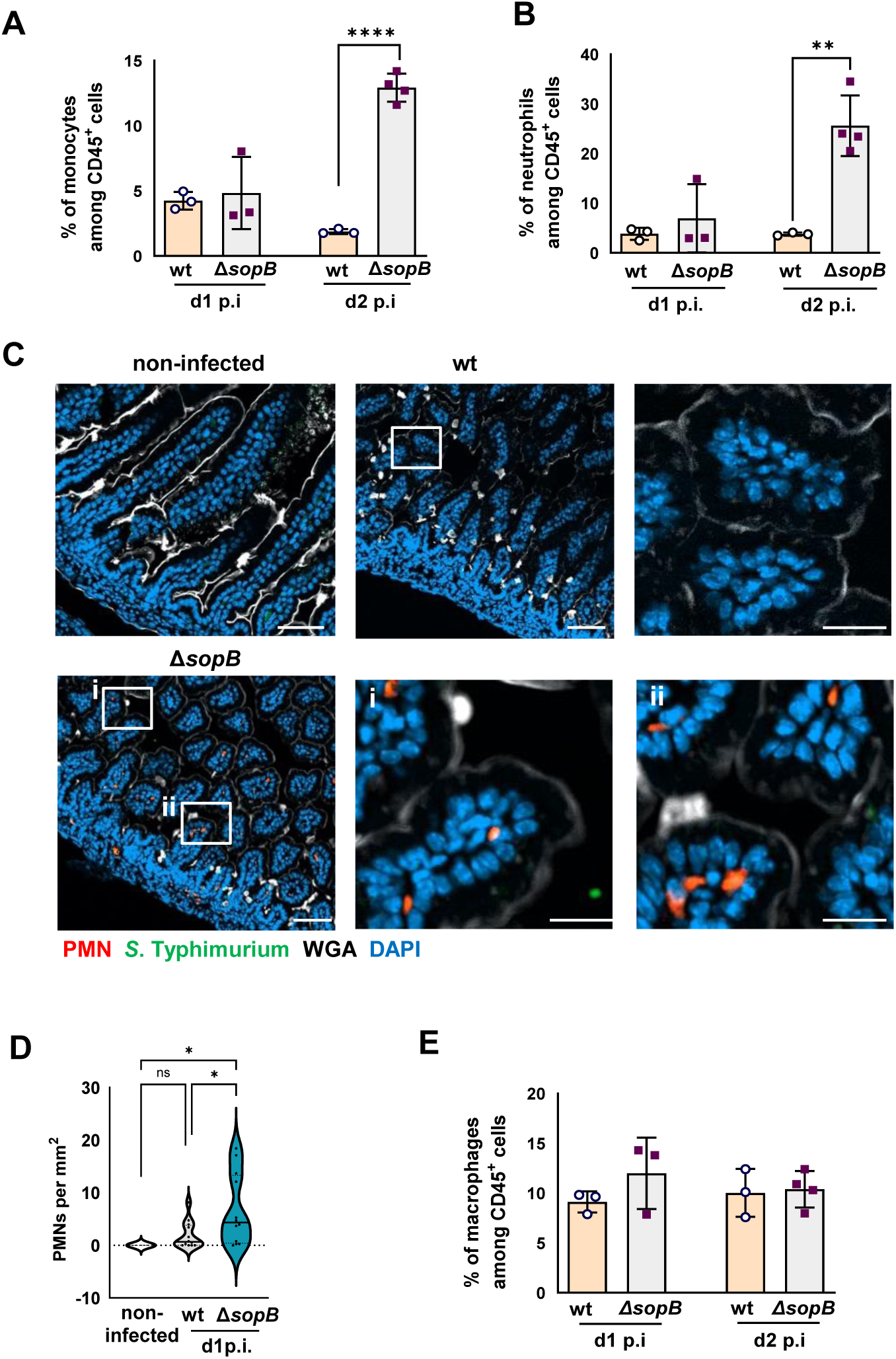
Immune cell infiltration in mucosal tissue. **(A-B)** 1-day-old C57BL/6 neonates were orally infected with 100 CFU wt (n=3 at day 1 and 2 p.i.) or *sopB* deficient (Δ*sopB*, n=3 and 4 at day 1 and 2 p.i., respectively). Flow cytometric analysis of *lamina propria* Ly6C^hi^Ly6G^-^CD11b^+^ MHCII^lo/-^CD45^+^DAPI^-^ monocytes **(A)** and Ly6G^+^Ly6C^int^CD11b^+^ MHCII^lo/-^CD45^+^DAPI^-^ neutrophils **(B). (C)** Immunostaining of neonatal small intestine tissue sections of age-matched uninfected animals (n=3) and neonate mice infected with wt (n=3) or *sopB* deficient (Δ*sopB*, n=3) *S*. Typhimurium (green) at day 1 p.i. for Ly6G^+^ neutrophils (red). Counter staining with WGA (white) and DAPI (blue). Bar=50 µm, white boxes indicate the enlarged panels to the right, insert i and ii: 20 µm. Representative images are shown. **(D)** Number of Ly6G^+^ neutrophils on 3-12 images with the total area of 3.8 mm^2^ of tissue sections of uninfected, age-matched neonates (n=3) or neonates infected with wt (n=4) or Δ*sopB* (n=4) *S*. Typhimurium at day 1 p.i.. One-way-ANOVA with multiple comparison test. **(E)** Flow cytometric analysis of *lamina propria* CD64^+^MHCII^+^CD45^+^DAPI^-^ macrophages at d 1 and d 2 p.i.. Mean ± SD, One-way-ANOVA with Bonferroni’s multiple comparison test. One data point represents one animal. ns, non-significant; *, p<0.05; **, p<0.01; ****, p<0.0001.

**Figure 4.**
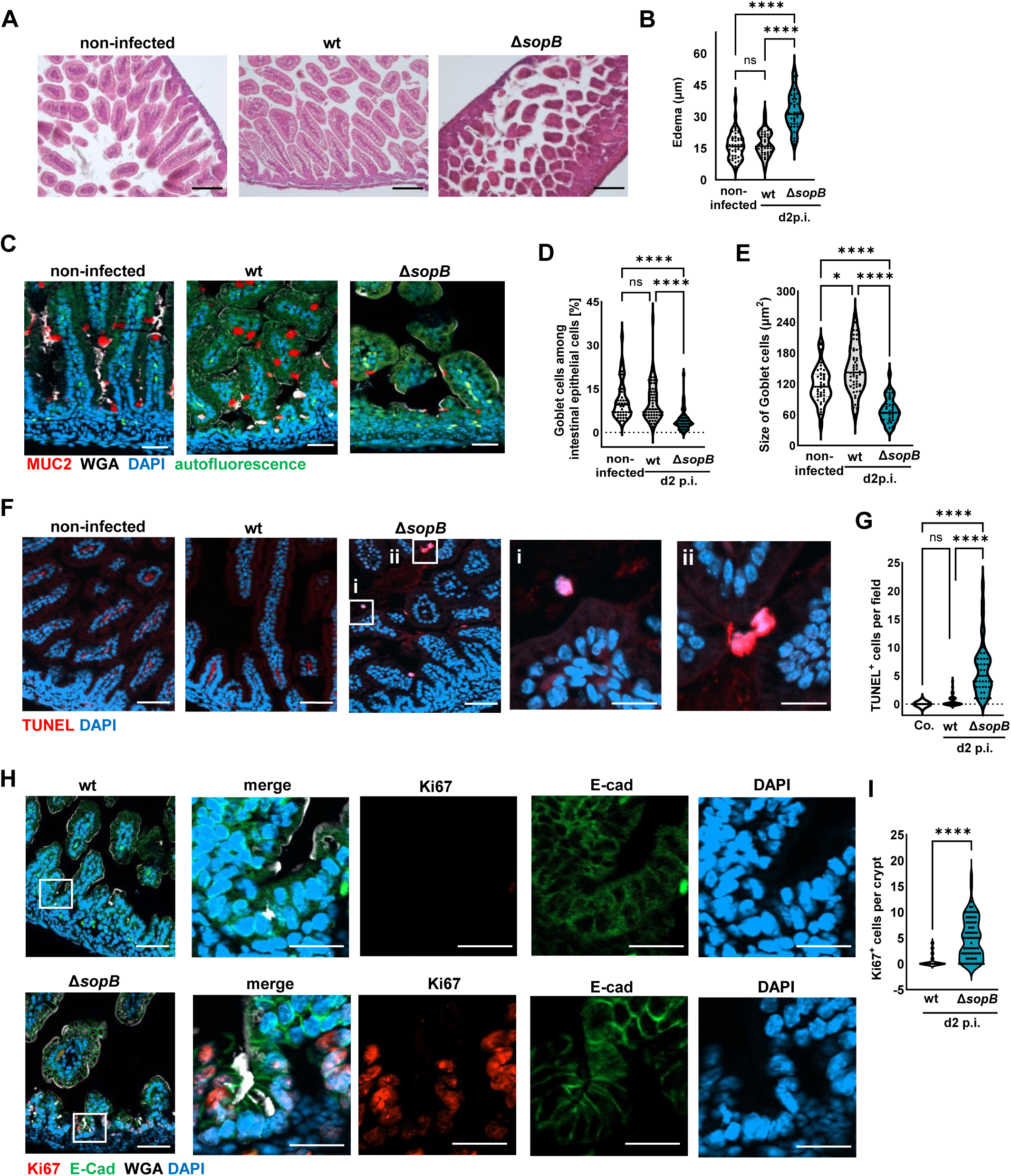
Histological characterization of the neonatal small intestine following infection. **(A)** Small intestinal tissue sections stained with H&E of non-infected (n=3) neonates or neonates infected with wt (n=3) or Δ*sopB* (n=3) *S*. Typhimurium at day 2 p.i.. Bar=100 µm. **(B)** Quantification of the depth of the *lamina propria* as a measure of the tissue edema measured in 15-20 different areas of one tissue section per animal (non-infected neonates (n=3) or neonates infected with wt (n=3) or Δ*sopB* (n=3)). **(C)** Muc2 immunostaining (red) in small intestine tissue sections of non-infected (n=3) neonates and the neonates infected with wt (n=3) or Δ*sopB* (n=3) *S*. Typhimurium at day 2 p.i.. Counterstaining with WGA (white), DAPI (blue); autofluorescence (green). Bar=50 µm. **(D)** Percentage of goblet cells among intestinal epithelial cells in non-infected (n=3) neonates and neonates infected with wt (n=3) or Δ*sopB* (n=3) *S*. Typhimurium at day 2 p.i.. 10-20 images with the size of 312µm x 250µm were evaluated on one tissue section per animal. **(E)** Goblet cell size in tissue sections of uninfected (n=3) neonates and neonates infected with wt (n=3) or Δ*sopB* (n=3) *S*. Typhimurium at day 2 p.i.. 11-21 goblet cells (µm^2^) were analyzed on each section. **(F)** TUNEL staining (red) of neonatal small intestine tissue sections of non-infected (n=2) neonates and neonates infected with wt (n=5) or Δ*sopB* (n=4) *S*. Typhimurium at day 2 p.i.. White boxes indicate enlarged images (i, ii). Counterstaining with DAPI (blue). Bar=50 µm; insert, 20 µm. **(G)** Number of TUNEL positive cells in tissue sections of uninfected (n=2) neonates and the neonates infected with wt (n=5) or Δ*sopB* (n=4) *S*. Typhimurium at day 2 p.i. 9 -12 images with the size of 624µm x 501µm were evaluated per animal. **(H)** Ki67 (red) immunostaining on small intestine tissue sections of the neonates infected with wt (n=3) or Δ*sopB* (n=4) *S*. Typhimurium at day 2 p.i.. Counterstaining with E-cadherin (green), WGA (white), and DAPI (blue). Bar=50 µm, insert: 20 µm. **(I)** Number of Ki67 positive cells in small intestinal tissue sections of uninfected (n=2) neonates and the neonates infected with wt (n=5) or Δ*sopB* (n=4) *S*. Typhimurium at day 2 p.i. 5-29 crypts were analyzed per section. Representative images are shown. Quantified data are shown as individual points in violin plots, solid lines represent the median. Data were analyzed using One-way ANOVA Kruskal–Wallis test with Dunn’s multiple comparisons post-test. ns, non-significant; *, p<0.05; ****, p<0.0001.

### TNF and necroptosis cause early disease progression and mortality

Inflammation-induced programmed cell death of epithelial cells may contribute to tissue inflammation through barrier disruption and promote disease progression after Δ*sopB S.* Typhimurium infection. Next, we tested mice impaired in different cell death pathways. The accelerated disease progression after Δ*sopB versus* wt *S.* Typhimurium infection was preserved in caspase 1 (Casp1) and ASC deficient mice (Fig. 5A-B) ^37^. Also, early mortality after Δ*sopB S.* Typhimurium infection was still observed in intestinal epithelial cell-specific caspase 8 deficient (*Casp8*^ΔIEC^) animals (Fig. 5C) and expression of the endogenous apoptosis regulator *Bcl2* or the nitric oxide generating protein iNOS (Nos2) remained unaltered in Δ*sopB S.* Typhimurium infected animals (Suppl. Fig. 4A-C) ^25,27^. These results make pyroptosis or apoptosis an unlikely explanation for early mortality after Δ*sopB S.* Typhimurium infection. In contrast, *Mlkl*^-/-^ animals infected with Δ*sopB S.* Typhimurium exhibited a significantly prolonged survival suggesting that necroptosis may contribute to the accelerated disease progression ^38^ (Fig. 5D). Consistently, histological analysis revealed reduced thickening of the submucosal tissue after Δ*sopB S.* Typhimurium infection of *Mlkl*^-/-^ mice (Fig. 5E and F).

**Figure 5.**
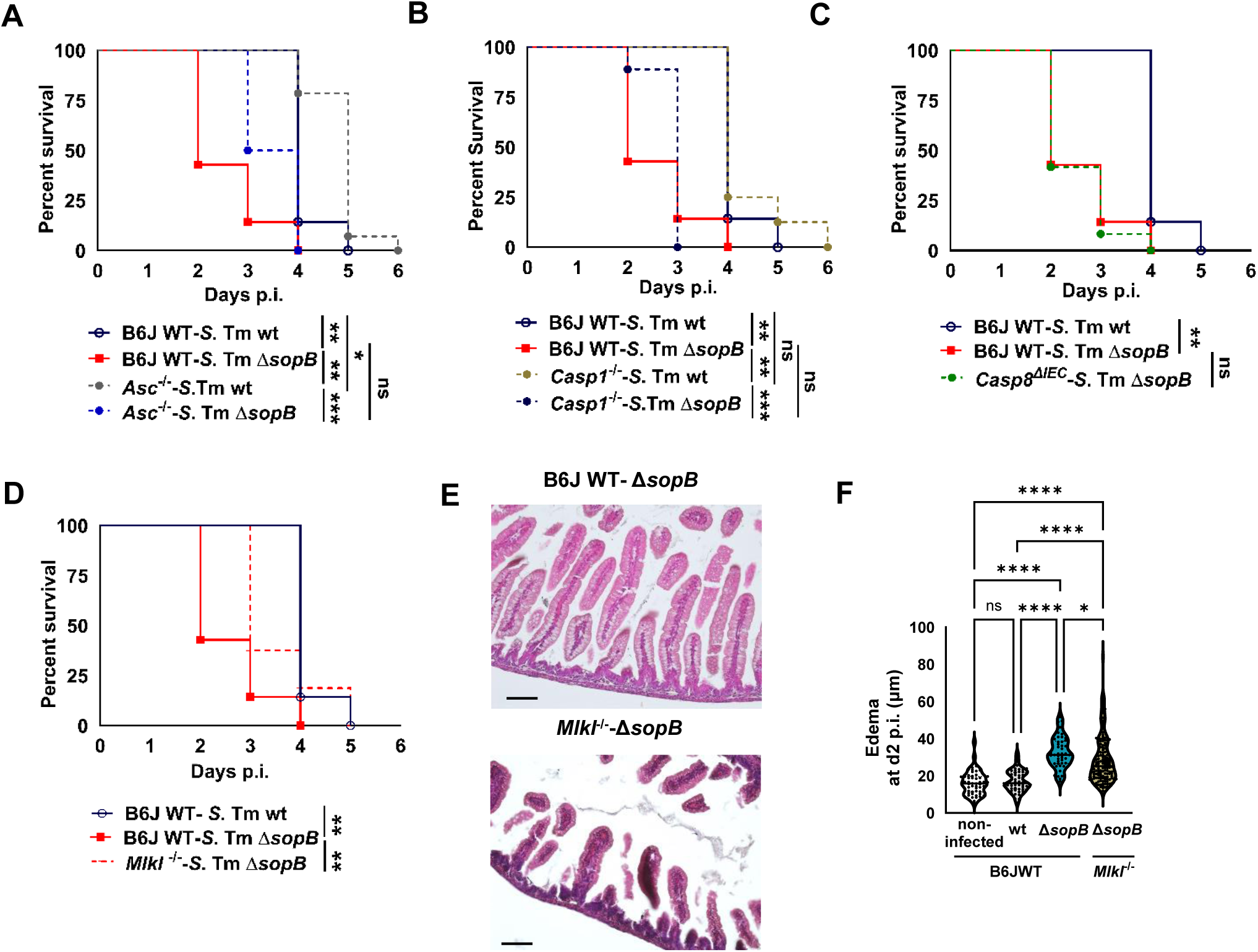
Cell death pathway contributing to the premature death of Δ*sopB S.* Typhimurium infected animals. **(A-D)** Kaplan-Meier curve of 1-day-old C57BL/6 wildtype (B6JWT, n=7 for both groups) and *Asc*-deficient (*Asc*^-/-^, n=6-14) **(A)**, wildtype and caspase 1 deficient (*Casp1*^-/-^, n=8-9) **(B)**, wildtype and intestinal epithelial cell-specific caspase 8 deficient (*Casp8*^ΔIEC^, n=6) **(C)**, and wildtype and Mlkl deficient (*Mlkl*^-/-^, n=16) **(D)** neonates orally infected with 100 CFU wt or *sopB* deficient (Δ*sopB*) *S.* Typhimurium. Note that the groups of wt and Δ*sopB S.* Typhimurium-infected wildtype mice are identical to Figure 1A. **(F)** H&E staining of small intestine tissue sections of B6JWT (n=3) and *Mlkl*^-/-^ (n=3) neonates infected with Δ*sopB S*. Typhimurium at day 2 p.i.. Bar=50 µm. log-rank (Mantel-Cox) test. **(G)** Quantification of the depth of the *lamina propria* as a measure of the tissue edema measured in 4-12 different areas of each tissue section of wildtype and *Mlkl*^-/-^ mice infected with wt or Δ*sopB S*. Typhimurium at day 2 p.i.. Quantified data are shown as individual points in violin plots, solid lines represent the median. Data were analyzed using One-way ANOVA Kruskal–Wallis test with Dunn’s multiple comparisons post-test. ns, non-significant; *, p<0.05; ****, p<0.0001.

Necroptosis is induced by TNF receptor (TNFR) stimulation prompting us to test TNF expression in intestinal tissue and the phenotype of *Tnfr1* deficient mice after *S.* Typhimurium infection. TNF mRNA expression was enhanced at day 1 and 2 p.i. in total small intestinal tissue (Fig. 6A and B) and at day 2 p.i. also in intestinal epithelial cells (Suppl. Fig. 5A and B). Early mortality after Δ*sopB S.* Typhimurium infection was completely blunted in the absence of TNFR1 (Fig. 6C) and Δ*sopB S.* Typhimurium infection-induced tissue thickening of the *lamina propria* at 2 days p.i. was abolished (Fig. 6D and E). Notably, the number of intraepithelial *S.* Typhimurium and the bacterial organ load in the mesenteric lymph node were not altered in the absence of TNFR1 or MLKL (Suppl. Fig. 5C and D). Although a moderate reduction in the bacterial organ load was noted in spleen and liver tissue of *Mlkl*^-/-^ mice, no significant difference was observed between wildtype and *Tnfr1*^-/-^ animals (Suppl. Fig. 5E and F). Together, these results support TNF-induced necroptosis of the intestinal epithelium as likely mechanism contributing to the accelerated disease progression after Δ*sopB S.* Typhimurium infection ^38^.

**Figure 6.**
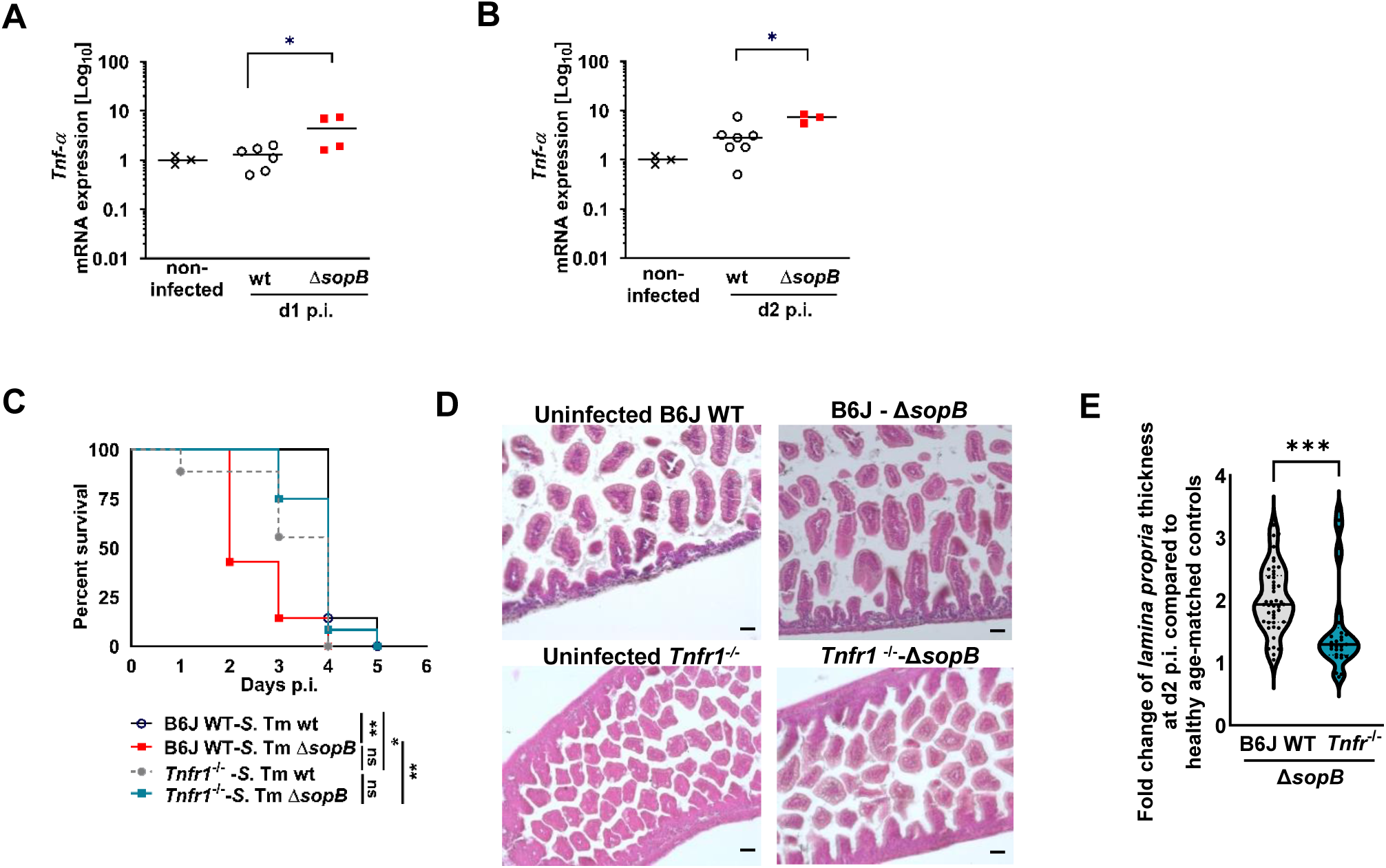
Role of TNF in neonate mice infected with Δ*sopB S.* Typhimurium. **(A and B)** mRNA expression level of *Tnf* in total gut tissue from age-matched non-infected animals (n=3) or neonates infected with wt (n=6-7) or Δ*sopB* (n=3-4) *S*. Typhimurium at day 1 **(A)** and day 2 **(B)** p.i.. Values were normalized to *Hprt* expression and are showed as the fold increase. Data were obtained from at least two independent experiments. One data point represents one animal, the median is indicated. **(C)** Kaplan-Meier curve of 1-day-old C57BL/6 wildtype (B6JWT, n=7-9) and *Tnfr1*-deficient (*Tnfr1*^-/-^, n=9-12) neonates orally infected with 100 CFU wt or *sopB* deficient (Δ*sopB*) *S.* Typhimurium. Note that the groups of wt and Δ*sopB S.* Typhimurium-infected wildtype mice are identical to Figure 1A. log-rank (Mantel-Cox) test. **(D)** H&E staining of small intestine tissue sections obtained at day 2 p.i.. from C57B/6 wildtype (n=3) and *Tnfr1*^-/-^ (n=5) neonate mice infected at day 1 after birth with Δ*sopB S*. Typhimurium. Bar=50 µm. **(E)** Fold change of the depth of the *lamina propria* as a measure of the tissue edema in small intestinal tissue sections of Δ*sopB S*. Typhimurium-infected over non-infected age-matched wildtype (n=3 and 3) and *Tnfr1*^-/-^ mice (n=3 and 3) at day 2 p.i. 3-9 different areas of the *lamina propria* on one tissue section per animal were analyzed. The quantified data are shown as individual points in violin plots, solid lines represent the median. Data were analyzed using Mann-Whitney Test. ns, non-significant; *, p<0.05; ***, p<0.001.

### S. Typhimurium-enterocyte interaction determines immunosuppressive effect of SopB

According to the current understanding of *S.* Typhimurium pathogenesis, the SPI1 effector SopB is translocated into the enterocyte together with other effector molecules that facilitate internalization and therefore specifically manipulates *Salmonella*-invaded enterocytes. However, *Salmonella*-invaded intestinal epithelial cells account for only approximately 1% of total enterocytes ^2^ which is in stark contrast to the strong epithelial chemokine expression and mucosal tissue TNF expression in the absence of SopB (Fig. 2 and Fig. 6). We therefore analysed whether SopB might also be secreted and act directly on epithelial and tissue immune cells. Consistent with a previous report on T3SS-dependent but translocon-independent SopB secretion ^28^, HA-tagged SopB was detected in conditioned cell culture medium and SopB secretion was dependent on the structural T3SS component InvC (Suppl. Fig. 6A). Expression and secretion of SopB were enhanced in absence of other SPI1 effectors. Also, treatment of TNF- and caspase inhibitor-exposed fibroblast L929 cells with recombinant SopB reduced the number of necroptotic cells indicating that secreted SopB may exert a biological function (Suppl. Fig. 6C-D). In addition, co-infection of a Δ*sopB S.* Typhimurium strain 1:1 with a SopB proficient, non-invasive triple mutant (Δ*sopEAsipA*) *S.* Typhimurium strain reduced early mortality suggesting that secreted SopB may counteract early inflammation induced by Δ*sopB S.* Typhimurium (Suppl. Fig. 6B) ^3^. However, co-infection of a Δ*sopB S.* Typhimurium strain 1:1 with a SopA but not SopB proficient non-invasive (Δ*sopEBsipA*) *S.* Typhimurium strain similarly inhibited early mortality (Suppl. Fig. 6B). Also, experiments using oral administration of recombinant SopB protein to neonatal mice infected with Δ*sopB S.* Typhimurium remained inconclusive. Thus, although we could confirm secretion and a cytoprotective effect of SopB *in vitro*, no *in vivo* evidence currently supports an extracellular activity of secreted SopB. Consistent with a solely direct effect of SopB on the chemokine expression of *S.* Typhimurium-invaded enterocytes, Δ*sopB S.* Typhimurium infection-induced accelerated disease progression and early mortality after were still observed in the absence of MyD88 signalling in hematopoietic cells such as immune cells (Fig. 7A). Also, immune cells such as neutrophilic granulocytes (neutrophils), monocytes and macrophages recruited to the intestinal mucosa of Δ*sopB versus* wt *S.* Typhimurium infected mice and sorted by flow cytometry, did not express higher levels of TNF mRNA after FACS sorting or TNF protein after intracellular cytokine staining (Fig. 7B-E, Suppl. Fig. 7 and 8A). The enhanced TNF mRNA levels in total intestinal tissue of Δ*sopB S.* Typhimurium infected mice (Fig. 6A and B) therefore most likely resulted not from increased expression but from enhanced recruitment of immune cells that constitutively express a certain level of TNF. Interestingly, the mean fluorescent intensity (MFI) of intracellular TNF staining was significantly lower in PMA/ionomycin re-stimulated monocytes from Δ*sopB* as compared wt *S.* Typhimurium-infected animals (Fig. 7F, Suppl. Fig. 8B). This reduction in intracellular TNF staining was only observed in monocytes and associated with a significantly reduced viability of these cells both under non-stimulated and PMA/ionomycin stimulated conditions (Fig. 7G-L, Suppl. Fig. 8C). Thus, the TNF-driven accelerated disease course may result from the inflammation-induced reduction in the viability of recruited monocytes and subsequent release of membrane-bound or intracellular TNF stores causing epithelial cell damage by necroptosis and loss of barrier integrity.

**Figure 7.**
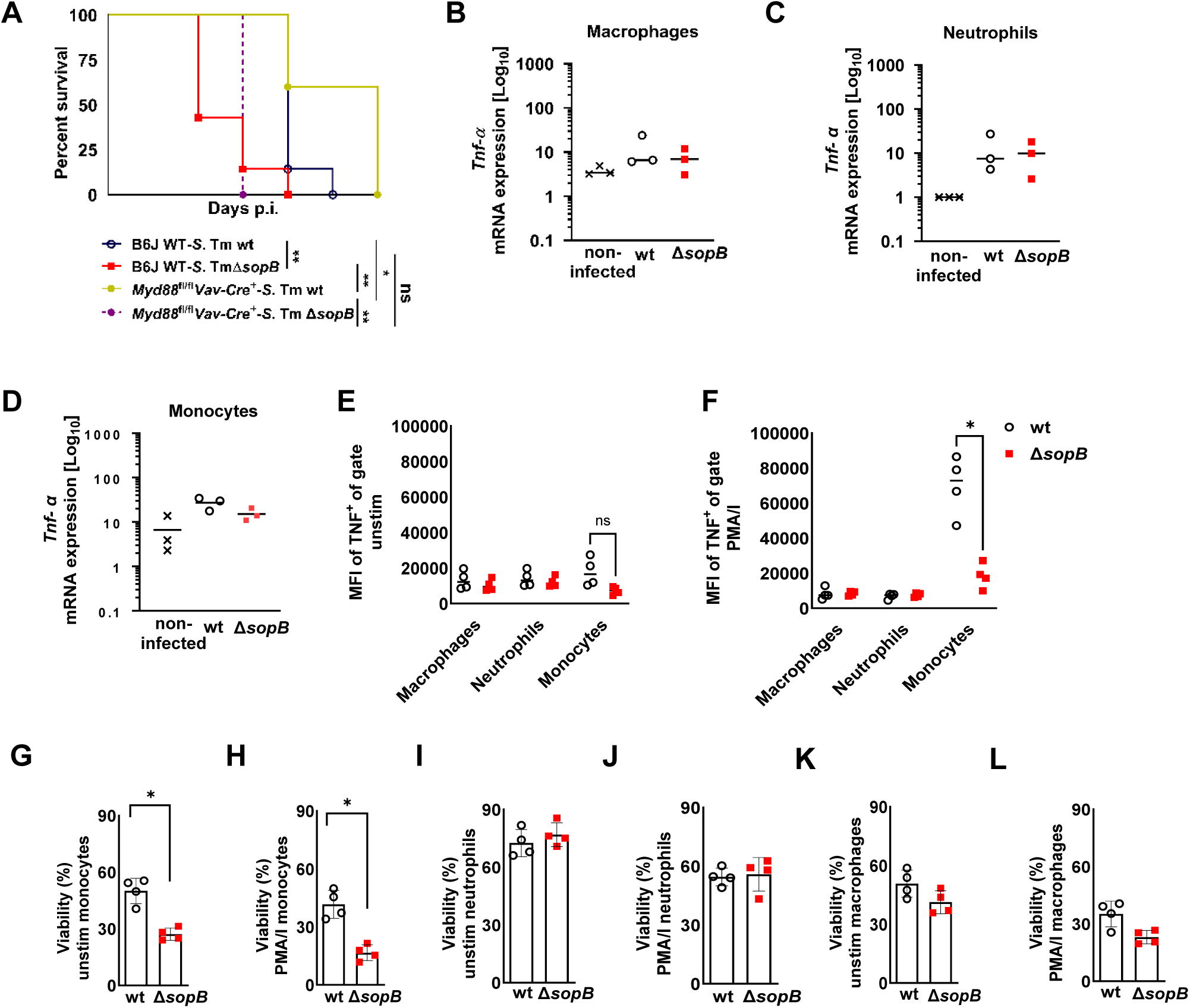
TNF staining in *lamina propria* immune cells of Δ*sopB S.* Typhimurium infected animals. **(A)** Kaplan-Meier curve of 1-day-old C57BL/6 wildtype (B6JWT, n=7) and hematopoietic cell-specific MyD88-deficient (*MyD8^fl/fl^VavCre^+^*, n=4-5) neonates orally infected with 100 CFU wt or *sopB* deficient (Δ*sopB*) *S.* Typhimurium. log-rank (Mantel-Cox) test. Note that the groups of wt and Δ*sopB S.* Typhimurium-infected wildtype mice are identical to Figure 1A. **(B-D)** Quantitative RT-PCR analysis of the *Tnf* mRNA expression in CD64^+^MHCII^+^CD45^+^DAPI^-^ macrophages **(B)**, Ly6G^+^Ly6C^int^CD11b^+^MHCII^lo/-^CD45^+^DAPI^-^ neutrophils **(C)**, and Ly6C^hi^Ly6G^-^CD11b^+^ MHCII^lo/-^CD45^+^DAPI^-^ monocytes **(D)** sorted by flow cytometry sorted from small intestinal *lamina propria* cells of non-infected mice (n=3), neonates infected with wt (n=3) or Δ*sopB* (n=3) *S*. Typhimurium at day 1 p.i.. Values were normalized to *Hprt* expression. Median, one-way ANOVA with Tukey’s multiple comparison test. **(E and F)** Mean fluorescence intensity (MFI) for TNF after intracellular cytokine staining of the indicated immune cell subsets isolated at day 2 p.i. from the *lamina propria* of wt (n=4) or Δ*sopB* (n=4) *S.* Typhimurium*-*infected animals without stimulation **(E)** or after stimulation with PMA/ionomycin (PMA/I). Median, two-way ANOVA with Sidak’s multiple comparison test. **(G-L)** Viability of unstimulated **(G, I, K)** or PMA/I stimulated **(H, J, L)** macrophages, neutrophils, and monocytes isolated from the *lamina propria* of wt (black circles, n=4) or Δ*sopB* (red squares, n=4) *S.* Typhimurium infected animals. Mean ± SD, Mann-Whitney test. ns, non-significant; *, p<0.05.

**Figure 8.**
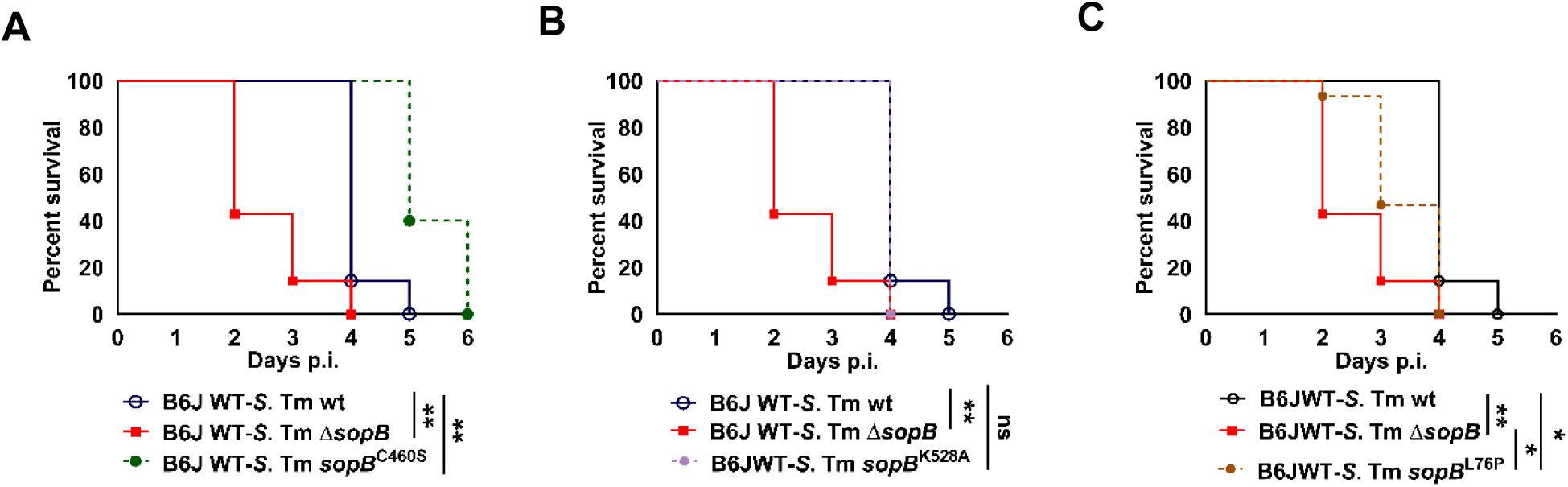
Survival of neonates infected with *S.* Typhimurium *sopB* mutants. **(A-C)** Kaplan-Meier curve of 1-day-old C57BL/6 wildtype mice orally infected with 100 CFU wt (n=7), *sopB* deficient (Δ*sopB,* n=7) *S.* Typhimurium, or S. Typhimurium strains carrying a point mutation in the C-terminal domain of SopB (*sopB*^C460S^, n=5; and *sopB*^K528A^, n=4) or N-terminal domain of SopB (*sopB*^L76P^, n=15). Note that the groups of wt and Δ*sopB S.* Typhimurium-infected wildtype mice are identical to Figure 1A. log-rank (Mantel-Cox) test. ns, non-significant; *, p<0.05; **, p<0.01.

### Phosphatidyl-inositol phosphatase independent influence of SopB

Point mutations at position 460 and 528 of SopB abolish its phosphatidyl-inositol phosphatase activity and associated downstream effects ^4,7,23^. Unexpectedly, mice infected with *S.* Typhimurium mutants with a chromosomal C460S or K528A mutation in SopB exhibited a wildtype-like or even protracted disease course and reduced mortality (Fig. 8A and B). In contrast, mice infected with *S.* Typhimurium carrying a chromosomal mutation in the N-terminal domain of SopB at position 76 (L76P) exhibited a significantly accelerated disease course and earlier mortality reminiscent of Δ*sopB S.* Typhimurium infected animals (Fig. 8C). SopB via its N-terminal CRIB like motiv (aa117-168) interacts with the small GTPase of the Rho family Cdc42 ^33,34^. The L76P point mutation was demonstrated to strongly impaire the interaction with Cdc42 but maintain the inositol phosphatidyl phosphatase activity ^32^. These results suggest that the observed accelerated disease course induced by Δ*sopB S.* Typhimurium is mediated by the lack of interaction of the N-terminal domain with host regulatory factors such as Cdc42 but not its phosphatidyl-inositol phosphatase activity.

## Discussion

Here we characterized the potent immunosuppressive effect of the *S.* Typhimurium SPI1 effector SopB in the neonatal infection model that dramatically retards tissue inflammation, reduces disease progression, and delays mortality *in vivo*. Although SopB in a number of studies has been shown to synergize with SopE/SopE_2_ and contribute to actin remodelling, ruffle formation, cell invasion, inhibition of lysosomal fusion, intracellular growth, and disease symptoms, our data suggest that its primary role is to suppress the antimicrobial immune response and delay disease progression ^3–5,7,9,10,12,13,15–17,19–21,39^. This is consistent with the previously described cell survival promoting activity of SopB via Akt stimulation, which, however, requires its phosphatidyl-inositol phosphatase activity ^22–24,26,28,33,40^. We show that this potent immune-evasive effect of SopB facilitates prolonged survival of the host and thereby most likely increased pathogen shedding enhancing the likelihood of host transmission.

SopB almost completely abolished the early epithelial chemokine expression at day 1 and immune cell recruitment at day 2 after oral infection of neonate mice. This is consistent with the early but also sustained detection of SopB and its activity in co-culture *in vitro* and in mucosal tissue *in vivo* ^24,25,29^. An important role of SopB for immune evasion is also consistent with the presence of the *sopB* gene in all clinical isolates ^8,38^. Notably, previous reports on the immunosuppressive role of SopB have revealed somewhat contradictory results. In the adult murine infection model, one study revealed no difference in the degree of colonic tissue edema and PMN infiltration following Δ*sopB* as compared to wt *S*. Typhimurium infection, whereas another study demonstrated enhanced colitis severity, goblet cell loss and bacterial translocation ^37,41^. Also, infection of bovine ileal loops with a Δ*sopB* mutant *S.* Dublin strain led to lower, not higher tissue infiltration of PMNs ^42^. In the neonatal mouse infection model, only the characterization of Δ*sopB S.* Typhimurium-infected animals allowed to detect the surprisingly early and potent epithelial response to infection with expression of chemokines, cytokines and the antimicrobial protein Reg3γ leading to enhanced immune cell recruitment and mucosal tissue inflammation. Given that animals in this model are infected with only 100 CFU orally and immune activation critically relies on a functional T3SS, *S.* Typhimurium must rapidly invade intestinal epithelial cells at significant numbers.

SopB has been well characterized as phosphatidyl-inositol phosphatase mediated by its C-terminal domain ^4,9,23^. This activity alters the phosphorylation status of phosphatidyl-inositol residues at the plasma membrane entry site to activate N-WASP and F-actin polymerization ^7,11,43,44^. Furthermore, the phosphatase activity has been linked to actin reorganization and invasion ^7,9,10^, membrane fission ^5^, inhibition of lysosomal fusion ^19^, ion flux alterations and fluid loss ^44^, as well as early Akt signaling, and host cell survival ^22–26,40^. However, two point mutations (C460S and K528A) within the C-terminal domain of SopB known to abolish the phosphatase activity had no influence on the course of the disease in neonatal mice making a contribution of the phosphatase activity to the observed phenotype unlikely ^4,7,23^. More recently, SopB was shown to interact with Cdc42 by the CRIB like motif in its N-terminal domain ^31,33,34^. Here it acts like a guanosine dissociation inhibitors (GDI) maintaining an inactive state of the small Rho GTPase Cdc42 ^34^. This interaction was absent when SopB carried a point mutation at position 76 (L76P) ^32^. Modification of the Cdc42 activity may directly or indirectly suppress early intestinal epithelial chemokine expression upon infection with SopB proficient *S.* Typhimurium. The molecular mechanism, however, remains to be elucidated ^45,46^.

Together, we demonstrate a pronounced delay in the disease progression and mortality mediated by SopB in the neonatal infection model *in vivo*. This activity was independent of the phosphatidyl-inositol phosphatase activity of the C-terminal domain of SopB but required an intact N-terminal domain. Mechanistically, SopB abolished early epithelial chemokine and cytokine expression, delayed immune cell infiltration, promoted the viability of infiltrating monocytes, and reduced the TNFR1-mediated effect on mucosal tissue integrity, enterocyte necroptosis, and host survival.

## Acknowledgement

We thank Regina Holland, Martina Leufgens, Simone Martin and Josefine Weber-Heynemann for technical support. This work was supported by the Flow Cytometry Facility and the Immunohistochemistry Facility, core facilities of the interdisciplinary Center for Clinical Research (IZKF) Aachen within the Faculty of Medicine at RWTH Aachen University. Financial support was provided by the Priority Program SPP2225 (project ID P446460159, Ho2236/18-1 project to M.W.H.), Collaborative Research Centers CRC1382 (project ID 403224013-SFB 1382 project B01, and A05 to M.W.H., M.v.B.) and CRC/TRR 359 (project ID 491676693-SFB/TRR359 project A01 to M.W.H.) from the German Research Foundation (DFG), and the ERC Advanced Grant EarlyLife (project ID: 101019157 to M.W.H.). Also, this research project was funded by the START-Program of the Faculty of Medicine at RWTH Aachen University.

---

**Suppl. Figure 1. (A)** Schematic diagram of the neonatal small intestinal epithelial stem cell organoid co-culture. Stem cell organoid cells grown as monolayers were infected for 1 h with wt, *sopB* deficient (Δ*sopB*), or *sopB* deficient *S*. Typhimurium complemented in trans with *sopB* and its chaperon *sigE* (Δ*sopB* p*sopBsigE*, compl.) followed by a 1h incubation in 100 µg/mL gentamicin at 37°C. Cells were washed three times, lysed in 0.1% triton and the viable bacterial number was analyzed by serial dilution and plating. **(B)** Invasion rate expressed as the number of detected intracellular bacteria relative to the number of administered bacteria × 100 (in %). One data point represents one infected well, mean ± SD, two independent experiments. One-way ANOVA with Tukey’s multiple comparison test. ****, p<0.0001.

**Suppl. Figure 2. (A and B)** 1-day-old C57BL/6 neonates were left uninfected (n=3-4) or orally infected with 100 CFU wt (n=5-11), *sopB* deficient (Δ*sopB*, n=4-14), or *sopB* deficient *S*. Typhimurium complemented in trans with *sopB* and its chaperon *sigE* (Δ*sopB* p*sopBsigE*, compl., n=4-8). *Reg3γ* and *Cxcl5* mRNA expression in total isolated intestinal epithelial cells at day 1 **(A)** and day 2 **(B)** p.i.. Values were normalized to the housekeeping gene *Hprt* and are showed as fold expression over uninfected age-matched control animals. One data point represents one animal from at least two independent experiments. Median, one way ANOVA with Kruskal-Wallis comparison test. **(C-O)** Data from the color-scaled heat map (z-score) shown in figure 2C. Serum concentration of the indicated cytokines and chemokines in pg/mL in the serum of mice infected at day 1 after birth with 100 CFU wt (n=3, empty black circles) or *sopB* deficient *S*. Typhimurium (Δ*sopB*, n=4, red circles) at day 1 and 2 p.i. Mean ± SD, two-way ANOVA. One data point represents one animal. ns, non-significant; *, p<0.05; **, p<0.01; ***, p<0.001; ****, p<0.0001.

**Suppl. Figure 3.** Gating strategy to identify *lamina propria* Ly6C^hi^Ly6G^-^CD11b^+^ MHCII^lo/-^ CD45^+^DAPI^-^ monocytes, Ly6G^+^Ly6C^int^CD11b^+^ MHCII^lo/-^CD45^+^DAPI^-^ neutrophils and CD64^+^MHCII^+^CD11b^+^CD45^+^DAPI^-^ macrophages by flow cytometric analysis.

**Suppl. Figure 4. (A and B)** Quantitative RT-PCR analysis for *Bcl2* mRNA expression in total intestinal epithelial cells isolated at day 1 **(A)** or 2 **(B)** p.i. from non-infected C57BL/6 neonates (n=3) or neonates infected at day 1 after birth with wt (n=2-6) or Δ*sopB* (n=5-6) *S*. Typhimurium. **(C)** *iNOS* mRNA expression in total intestinal epithelial cells isolated at day 2 p.i. from non-infected C57BL/6 neonates (n=5) or neonates infected at day 1 after birth with wt (n=7) or Δ*sopB* (n=8) *S*. Typhimurium. Median, one-way ANOVA with Tukey’s multiple comparison test. One data point represents one animal. ns, non-significant.

**Suppl. Figure 5. (A and B)** *Tnf-α* mRNA expression in total small intestinal epithelial cells isolated from non-infected age-matched animals (n=3) or neonates infected with wt (n=4-7) or Δ*sopB* (n=3-9) *S*. Typhimurium at day 1 **(A)** and **(B)** 2 p.i.. Values were normalized to *Hprt* expression and are showed as the fold increase over uninfected age-matched control animals. **(C-F)** Organ counts of *sopB* deficient (Δ*sopB*) *S*. Typhimurium in total isolated gentamicin treated intestinal epithelial cells **(C)**, mesenteric lymph nodes (MLN) **(D)**, spleen **(E),** and liver tissue homogenates **(F)** of C57BL/6 wildtype (B6JWT, n=3), *Tnfr*^-/-^ (n=6), and *Mlkl*^-/-^ (n=3) mice at day 2 p.i.. Median, one-way ANOVA with Tukey’s multiple comparison test. One data point represents one animal. *, p<0.05; **, p<0.01.

**Suppl. Figure 6. (A)** Western blot of HA-tagged SopB (65.1 KDa) in culture supernatant (supernatant) or whole bacterial lysates (pellet) using an anti-HA antibody. wt and *sopB* deficient (Δ*sopB*) *S.* Typhimurium or wt *S.* Typhimurium carrying an expression plasmid with HA-tagged SopB (wt p*sopB*::HA), a *sopA*, *sopE_2_*, *sipA* triple mutant encoding a *sopB*::HA-fusion protein (Δ*sopAE_2_sipA*, *sopB*::HA), a *sopA*, *sopE_2_*, *sipA,* and *sopB* quadruple mutant carrying an expression plasmid with HA-tagged SopB (Δ*sopABE_2_sipA*, p*sopB*::HA), or a *invC* deficient strain carrying an expression plasmid with HA-tagged SopB (Δ*invC,* p*sopB*::HA) were analyzed. **(B)** Kaplan-Meier curve of 1-day-old C57BL/6 (B6JWT) neonates orally infected with 100 CFU wt (n=10), *sopB* deficient (Δ*sopB*, n=20), or a 1:1 ratio of *sopB* deficient (Δ*sopB*) and *sopB, sopE_2_* and *sipA* deficient (Δ*sopBE_2_sipA*) (n=11) or of *sopB* deficient (Δ*sopB*) and *sopA, sopE_2_* and *sipA* deficient (Δ*sopAE_2_sipA*) (n=17) *S*. Typhimurium. log-rank (Mantel-Cox) test. **(C)** L929 cells were pretreated with 5 µM zVAD alone or together with 3 µg/mL SopB for 1 h, followed by treatment with 10 ng/mL TNF for 20 h. Left panel: representative microscopic images of treated L929 cells. The white arrow heads point to cells with a necrosis-like phenotype. Bar=20 µm. Right panel: Flow cytometric analysis of the number of AV^+^/PI^+^ necroptotic L929 cells by staining with Annexin V (AV) and propidium iodide (PI). The results are presented as mean ± SD. **(D)** L929 cells were pretreated with 10 µg/mL cycloheximide with or without 10 µM Nec-1s, 5 µM zVAD or 0.1 µg/mL SopB for 1 h, followed by treatment with 10 ng/mL TNF for 15 h. Left panel: representative microscopic images of treated L929 cells. Bar=20 µm, Right panel: Flow cytometric analysis of the number of AV^+^/PI^-^ apoptotic L929 cells. The results are presented as mean ± SD. One-way ANOVA with Tukey’s multiple comparison test. ns, non-significant; *, p<0.05; **, p<0.01; ***, p<0.001; ****, p<0.0001.

**Suppl. Figure 7.** Gating strategy to identify *lamina propria* Ly6C^hi^Ly6G^-^CD11b^+^ MHCII^lo/-^ CD45^+^DAPI^-^ monocytes, Ly6G^+^Ly6C^int^CD11b^+^ MHCII^lo/-^CD45^+^DAPI^-^ neutrophils and CD64^+^MHCII^+^CD45^+^DAPI^-^ macrophages by flow cytometric analysis.

**Suppl. Figure 8. (A)** Mean fluorescence intensity (MFI) for TNF after intracellular cytokine staining of CD45^+^ immune cells isolated at day 2 p.i. from the *lamina propria* of wt (n=4) or Δ*sopB* (n=4) *S.* Typhimurium*-*infected animals without stimulation or after stimulation with PMA/ionomycin (PMA/I). Median, two-way ANOVA with Sidak’s multiple comparison test. One data point represents one animal. **(B)** Flow cytometric staining of intracellular TNF in Ly6C^hi^Ly6G^-^CD11b^+^ MHCII^lo/-^CD45^+^DAPI^-^ monocytes, Ly6G^+^Ly6C^int^CD11b^+^ MHCII^lo/-^ CD45^+^DAPI^-^ neutrophils, and CD64^+^MHCII^+^CD45^+^DAPI^-^ macrophages isolated from wt (n=4) or Δ*sopB* (n=4) *S*. Typhimurium-infected mice at day 2 p.i. after re-stimulation with PMA/I. FMO, fluorescence minus one, TNF staining control; MFI, mean fluorescence intensity. **(C)** Flow cytometric analysis of the viability of PMA/I stimulated monocytes isolated from the *lamina propria* of wt (n=4) or Δ*sopB* (n=4) *S.* Typhimurium infected animals at day 2 p.i..

