## Supplemental figures for "Phosphatase-independent suppression of mucosal inflammation and disease progression by *Salmonella* SopB" (from another version of this preprint: version 1, 2024-09-02)

### Supplemental Figure 1

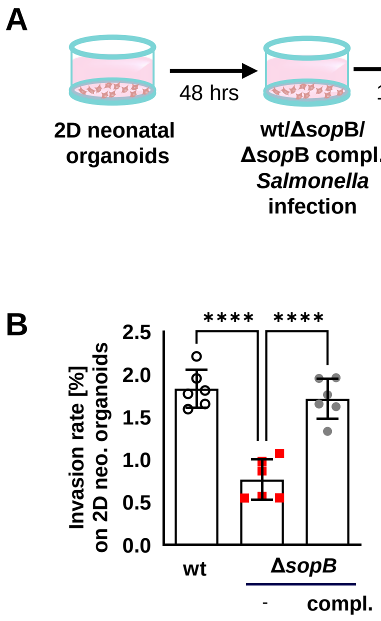

#### Supplemental Figure 2

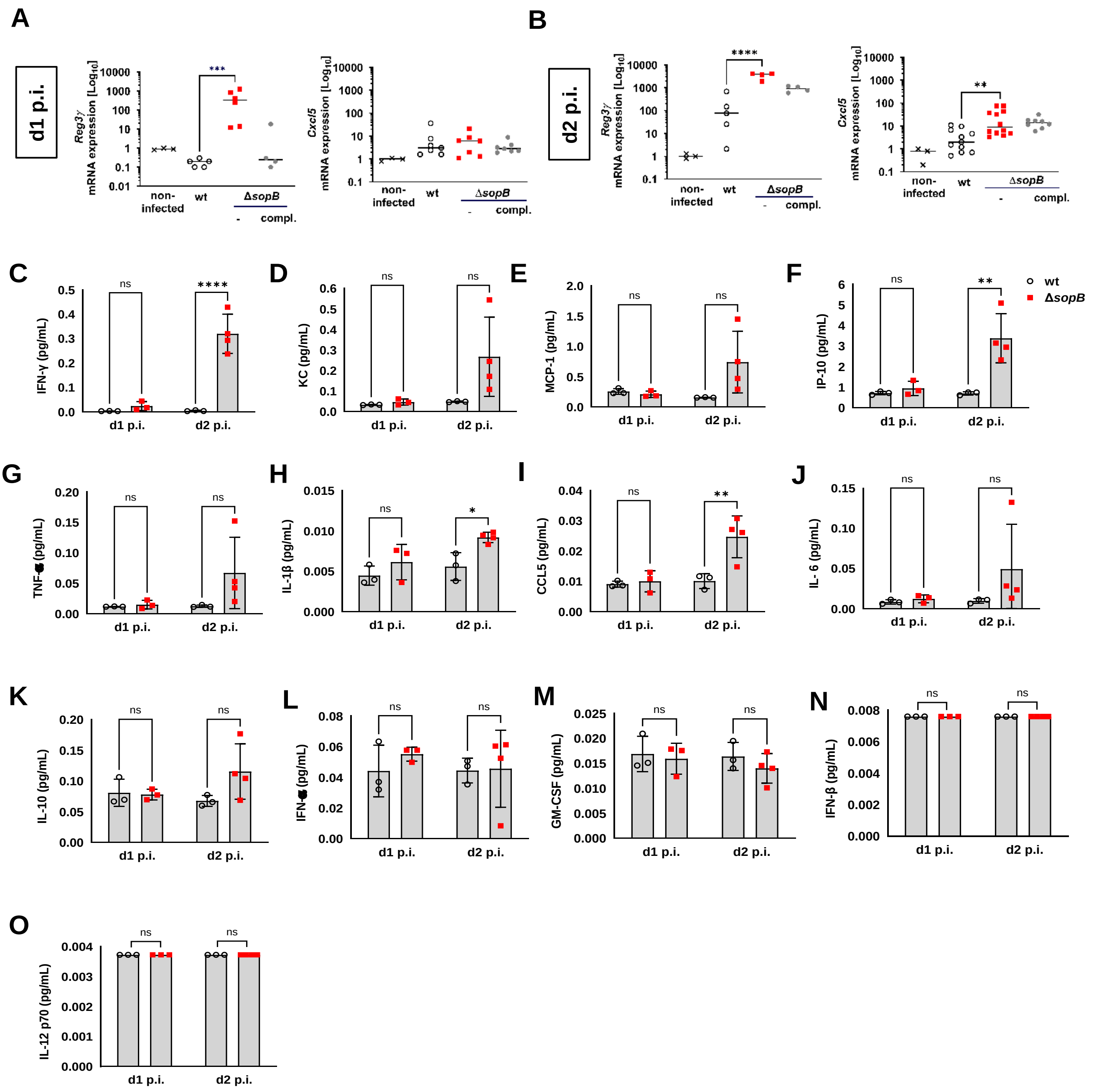

### Supplemental Figure 3

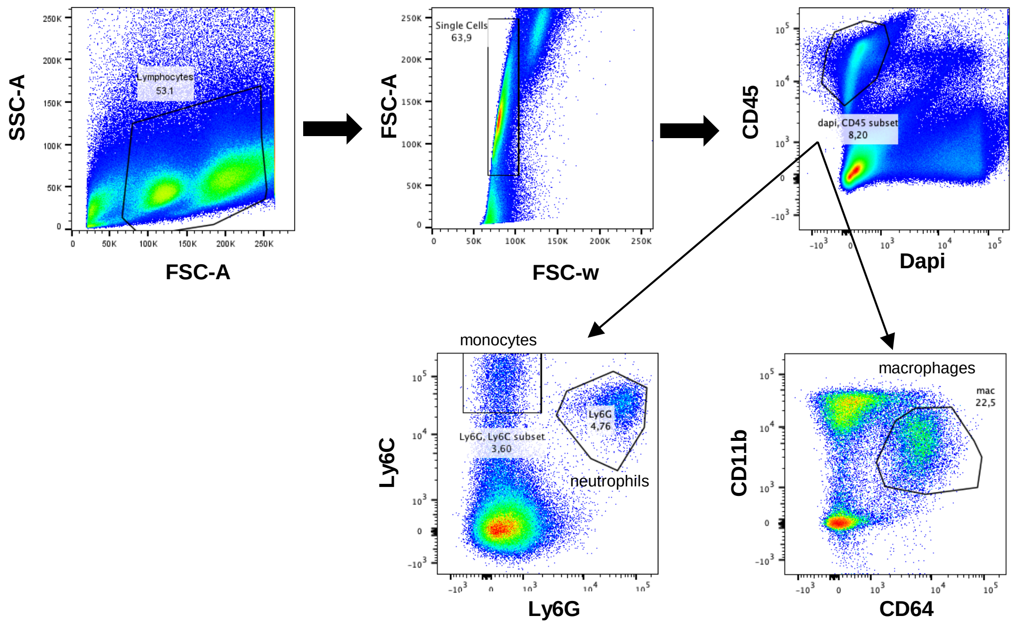

#### Supplemental Figure 4

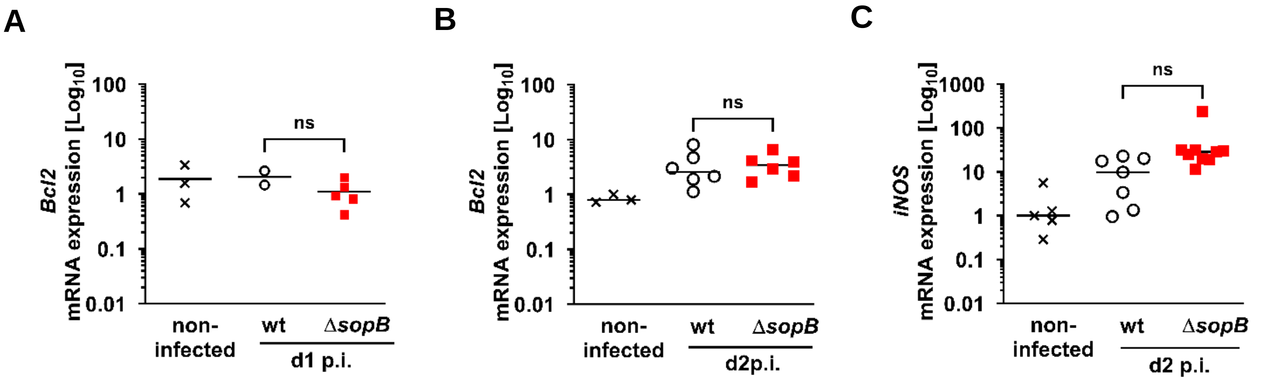

**Supplemental Figure 5**

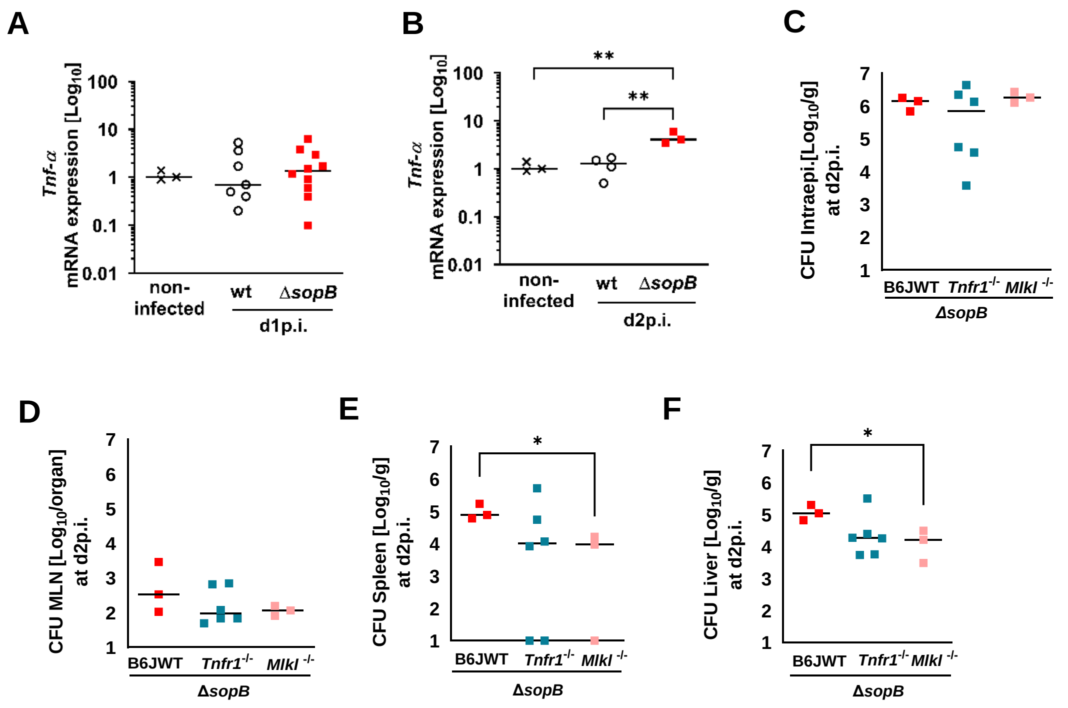

### Supplemental Figure 6

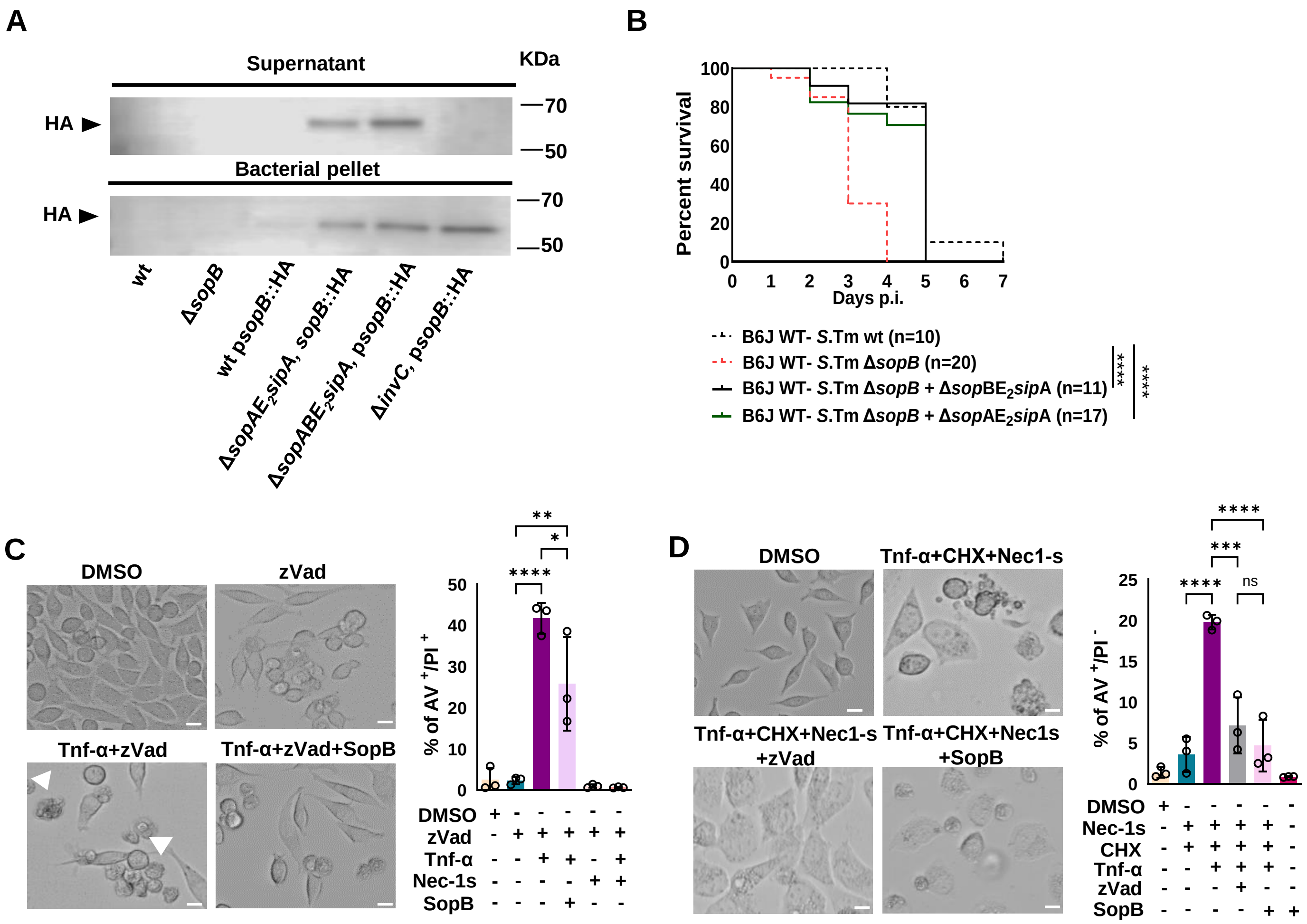

#### Supplemental Figure 7

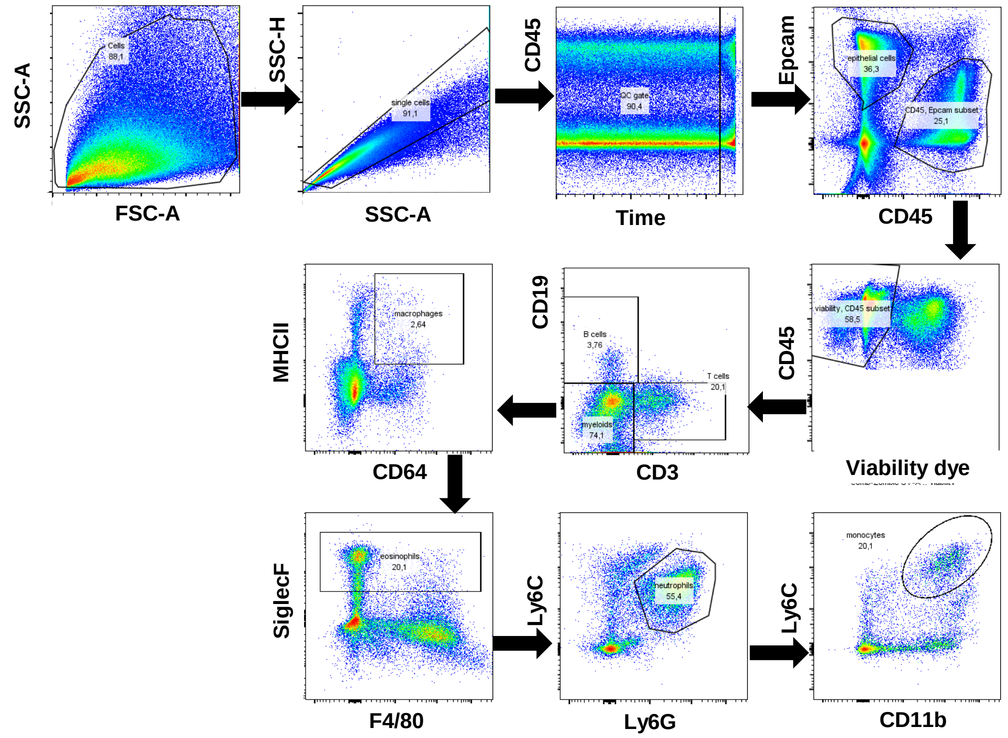

### Supplemental Figure 8

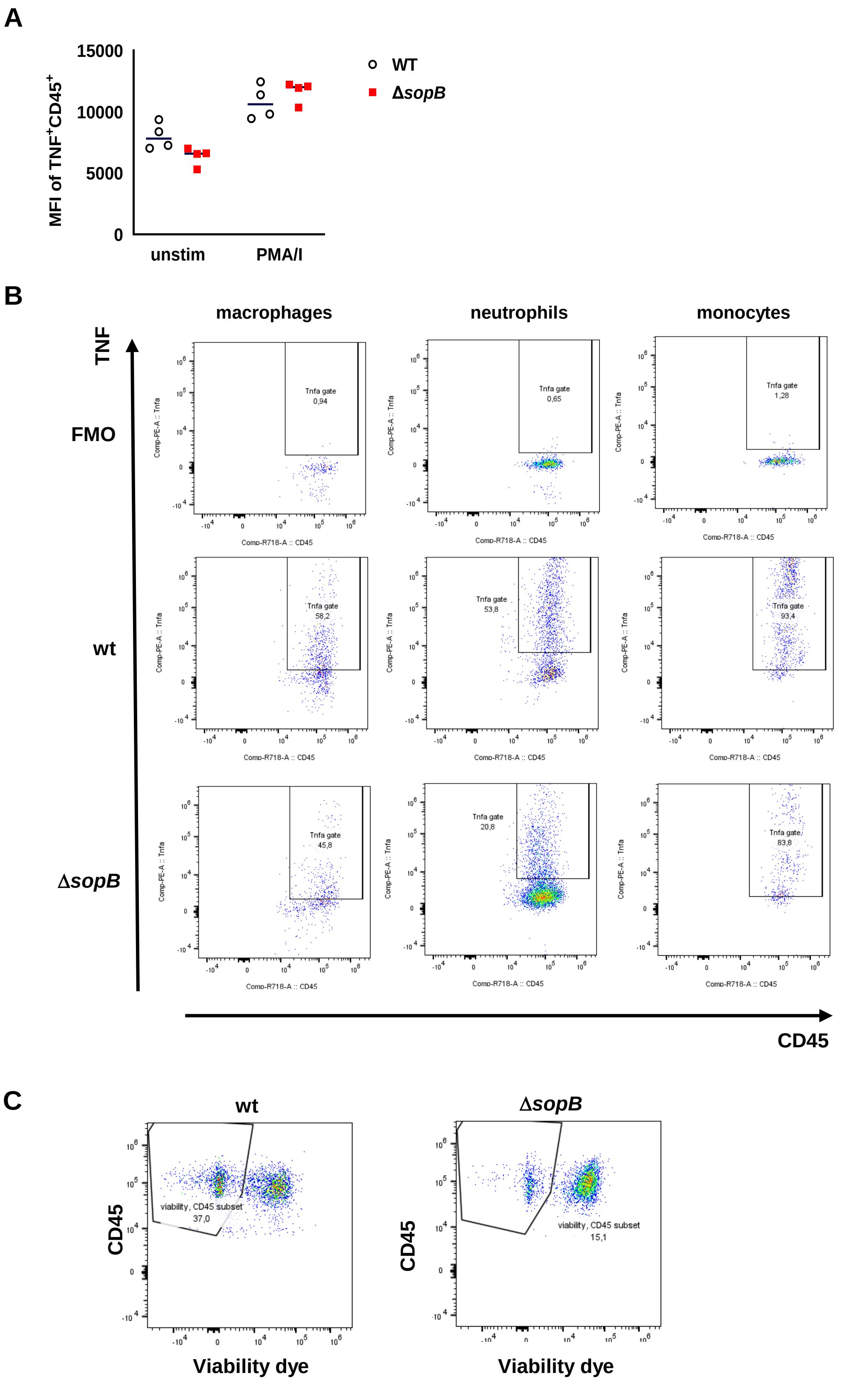
